# MINERVA FAIR assessment fosters open science & scientific crowd-sourcing in systems biomedicine

**DOI:** 10.1101/2024.08.28.610042

**Authors:** Irina Balaur, Danielle Welter, Adrien Rougny, Esther Thea Inau, Alexander Mazein, Soumyabrata Ghosh, Reinhard Schneider, Dagmar Waltemath, Marek Ostaszewski, Venkata Satagopam

## Abstract

The Disease Maps Project (https://disease-maps.org) focuses on the development of disease-specific comprehensive structured knowledge repositories supporting translational medicine research. These disease maps require continuous interdisciplinary collaboration and should be reusable and interoperable. Adhering to the Findable, Accessible, Interoperable and Reusable (FAIR) principles enhances the utility of such digital assets.

We used the RDA FAIR Data Maturity Model and assessed the FAIRness of the Molecular Interaction NEtwoRk VisuAlization (MINERVA) Platform. MINERVA is a standalone webserver that allows users to manage, explore and analyse disease maps and their related data manually or programmatically. We exemplify the FAIR assessment on the Parkinson’s Disease Map (PD map) and the COVID-19 Disease Map, which are large-scale projects under the umbrella of the Disease Maps Project, aiming to investigate molecular mechanisms of the Parkinson’s disease and SARS-CoV-2 infection, respectively.

We discuss the FAIR features supported by the MINERVA Platform and we outline steps to further improve the MINERVA FAIRness and to better connect this resource to other ongoing scientific initiatives supporting FAIR in computational systems biomedicine.

## 2 Introduction

The FAIR principles - Findability, Accessibility, Interoperability and Reusability - are guidelines towards improving data reusability, focusing on machine-assisted data handling for the minimization of human error^1^. Implementation of such guidelines is an important and challenging task, supported by numerous international initiatives. One of them is the Research Data Alliance (RDA: https://www.rdalliance.org/), a community-driven effort for supporting development of infrastructures for data sharing and re-use^2, 3^. Alignment with and implementation of the FAIR principles is a core aspect of many RDA initiatives and projects, such as the FAIR Data Maturity Model Working Group^4^. Elsewhere, the Innovative Medicines Initiative (IMI) FAIRplus project (https://fairplus-project.eu/) is a scientific academia-industry collaboration focusing on the development of tools and guidelines for the FAIRification in translational medicine projects such as the FAIR Cookbook^5^.

In systems biomedicine, scientific communities continuously integrate the FAIR principles, for example, the German National Research Data Infrastructure (https://www.dfg.de/en/researchfunding/funding-initiative/nfdi). On international scale, the umbrella Disease Maps Project (https://disease-maps.org/)^6, 7, 8^ and its various projects, including the Parkinson’s Disease Map (PDmap: https://disease-maps.org/parkinsons/)^9, 10, 11^ and the COVID-19 Disease Map (https://covid.pages.uni.lu/) ^12, 13, 14^, implement FAIR principles to increase the value of scientific outcomes. The PD map and the COVID-19 Disease Map are sets of diagrams describing molecular mechanisms involved in the Parkinson’s disease^9, 10^ and SARS-CoV-2 infection^12, 13, 14^, respectively. They feature manually curated and annotated information extracted from the relevant literature. These resources are developed and supported by active communities^9, 10, 12^ and can be accessed through the Molecular Interaction NEtwoRk VisuAlization (MINERVA) Platform^15^ as follows: PD map at https://pdmap.uni.lu/minerva/ and COVID-19 Disease Map at https://covid19map.elixir-luxembourg.org/minerva/

MINERVA is a platform for the visualization and curation of systems biology diagrams and provides several functionalities for their analysis, such as the automatic annotation of their content, data overlay and conversion between different systems biology formats. MINERVA-hosted diagrams support in-depth analysis of disease-related pathways, and of their perturbations. The PD map and the COVID-19 Disease Map have been used as entry points for molecular diagnostics or identification of potential new and repurposed drug targets in PD and COVID-19 diseases, respectively, using various analysis techniques and pipelines, e.g. data integration, omics analysis, computational modelling, text mining and AI)^10, 11, 14^. Such analyses, however, require an easy access and reuse of maps and of their content.

We ran a detailed FAIR assessment of the PD map and of the COVID-19 Disease Map in the MINERVA Platform to evaluate how Disease Maps can be accessed and reused. We present this assessment and highlight the challenges inherent to the FAIRification of biomedical content in diagrammatic and network format rather than in tabular format. We emphasize that this assessment reflects the FAIR features supported by MINERVA (infrastructure) rather than the specific FAIRness of the individual resources (maps). We conclude that any FAIR improvement in MINERVA will be propagated across the entire Disease Maps community. To the best of our knowledge, this is among the first scientific efforts on performing the FAIR assessment of system biology structured computational resources (with data and metadata encoded in a graph-based diagram).

## 3 Results

We assessed the MINERVA FAIRness using the PDmap (https://pdmap.uni.lu/minerva/) and COVID-19 Disease Map (https://covid19map.elixir-luxembourg.org/minerva). The MINERVA FAIR score is 82.05% when all indicators are considered. More specifically, the Essential indicators scored 73,68% and the remaining indicators (with Important and Useful priorities) scored 90%. The detailed FAIR assessment of MINERVA using the FAIRplus indicator template is available at https://doi.org/10.5281/zenodo.10910033^16^.

In summary, the MINERVA Platform is a FAIR-compliant infrastructure:

- Findability: Facilitates the retrieval of biological maps via globally unique and persistent identifiers (URLs) for resources, alongside an indexing system that enables the exploration of integrated content.
- Accessibility: Ensures rapid Web-based access to the biological map content without the need for specialized software, tools, or settings. It also allows for access to the content via REST and JavaScript-based APIs (more details at https://minerva.pages.uni.lu/doc/api/) and implements secure authentication and authorization protocols upon project request.
- Interoperability: Offers compatibility with major systems biology standards, including SBML, CellDesigner, SBGN v03, GPML^21^, and BioPAX ^22^ through an integrated converter.
- Reusability: Provides flexible licensing options at the diagram or project level.

## 4 Discussion

As indicated in Table 1, the RDA scoring shows that most of the FAIR principles (https://www.go-fair.org/fair-principles/) are fulfilled. Interestingly, several FAIR RDA indicators were not applicable because the modularity of data and metadata are missing. Specifically, the representation of the model contains data (molecular entities and their relationships) and metadata (annotations of the biological content) within the same object. Thus, the Accessibility Essential indicator of “RDA-A2-01M: Metadata is guaranteed to remain available after data is no longer available” was not applicable since the removal of an object representing the molecular entity/ relationship (data) from the diagram, also implies deletion of its annotation (metadata) in the data representation model in MINERVA (details in Methods).

**Table 1.** Assessment of the FAIR principles in MINERVA (the Map)

| Principles | Status of the principle during the assessment | Explanations and comments |
| --- | --- | --- |
| F1. (Meta)data are assigned a globally unique and persistent identifier | Fulfilled at the level of data only; ongoing work for metadata | Data (maps and integrated biological content) is available online at https://pdmap.uni.lu/minerva/ and https://covid19map.elixir-luxembourg.org/minerva/ for PD map and COVID-19 Disease Map, respectively. For metadata, we started communication with the MINERVA development team on options to provide an additional store system for metadata (annotations). The MINERVA-Net is a registry to allow users to share information about their disease maps available in MINERVA, including the PD map and COVID-19 Disease Map, and to store these annotations in its centralized server<sup>17</sup>. |
| F2. Data are described with rich metadata (defined by R1 below) | Fulfilled | The content of maps is connected to well-established systems biomedicine resources (including UniProt, ChEMBL etc.) as well as is annotated with information regarding the initial publications supporting the integrated relationships/ knowledge. |
| F3. Metadata clearly and explicitly include the identifier of the data they describe | Not applicable | Given the data representation model in MINERVA with the data and metadata encoded in a single object (see Methods), the metadata does not refer to the data, but it is incorporated in the data. |
| F4. (Meta)data are registered or indexed in a searchable resource | Fulfilled | Search interface is implemented in MINERVA to allow search by name, identifier, reference or PubMed ID to identify elements and interactions. Disease maps (including the PD map and the COVID-19 Disease Map) |
|  |  | are indexed in the MINERVA-Net registry (https://minerva-net.lcsb.uni.lu)<sup>17</sup>. |
| A1. (Meta)data are retrievable by their identifier using a standardised communications protocol | Fulfilled at the level of data only; ongoing work for metadata (at the moment of the FAIR assessment) | The data and metadata can be accessed both manually (via online access and visual exploration in MINERVA) and automatically (via the MINERVA REST API). Moreover, data resolves to digital objects; for example, the biological maps are given by the MINERVA diagram objects, the biological entities are represented as objects of Protein, Metabolite etc. and their inter-relationships by connecting edges (arcs). However, the metadata does not contain information regarding access to data since metadata is incorporated in the data object (the data representation, see Methods). Finally, we assessed that currently the metadata identifier does not resolve to a metadata record (the RDA-A1-03M indicator). |
| A1.1 The protocol is open, free, and universally implementable | Fulfilled | The content of the Map is accessible online via the HTTPS protocol. |
| A1.2 The protocol allows for an authentication and authorisation procedure, where necessary | Fulfilled | The MINERVA Platform provides a uniform API for machine-access to the biological content integrated in the maps as well as it supports authentication and authorization procedures where requested. |
| A2. Metadata are accessible, even when the data are no longer available | Not applicable | Given the data representation model in MINERVA where the data and metadata are encoded in a single object, removing the data (the map or some molecular entities and their inter-relationships) will delete the metadata (associated annotations). |
| I1. (Meta)data use a formal, accessible, | Fulfilled | The content of the Map follows the CellDesigner SBML format that is an XML-based systems biology |
| shared, and broadly applicable language for knowledge representation. |  | standard<sup>18</sup>. MINERVA also provides functionality for conversion to other standard formats<sup>15</sup>, <sup>19</sup>. |
| **I2.** (Meta)data use vocabularies that follow FAIR principles | Fulfilled | The content of the Map is connected to biological resources (e.g. UniProt, ChEML etc.) and has annotations regarding the evidence provenience from initial publications. Ongoing efforts are on connecting the MINERVA integrated content to the Bricks Ontology e.g. <sup>20</sup>, <sup>14</sup>. |
| **I3.** (Meta)data include qualified references to other (meta)data | Fulfilled | The data includes qualified references to other data given that the biological content describes the molecular mechanisms of the Parkinson's disease and COVID-19, respectively and integrates the relationships among the various biological entities involved in these diseases (such as proteins involved in transport and translocation processes etc.). |
| **R1.** (Meta)data are richly described with a plurality of accurate and relevant attributes | Fulfilled | The SBML data model allows assignment of multiple stable identifiers to elements, interactions, and disease maps to identify their function and scope. The MINERVA Platform handles a broad range of stable identifiers<br>[(https://minerva.pages.uni.lu/doc/release_notes/v16.1/supported-annotations/)](https://minerva.pages.uni.lu/doc/release_notes/v16.1/supported-annotations/). |
| **R1.1.** (Meta)data are released with a clear and accessible data usage license | Not fulfilled from the indicators at the moment of the initial assessment; currently, available externally in e.g. FAIRDOMHub and in MINERVA v17.1 | The content of the Map is accessible under the CC-BY 4.0 usage licence <sup>13</sup>. During assessment the licensing information was available publicly on FAIRDOMHub, separately for each diagram of the COVID-19 Disease Map, e.g. https://fairdomhub.org/models/790. Now MINERVA v17.1 supports explicit definition of licenses for hosted maps. |
| **R1.2.** (Meta)data are | Fulfilled | The diagrams of the Map are annotated with informa- |
| associated with detailed provenance |  | tion on initial publications for evidence provenience (PubMed identifiers). |
| **R1.3.** (Meta)data meet domain-relevant community standards | Fulfilled | The diagrams of the Map are annotated also with systems biology standard identifiers for molecular entities including UniProt identifiers for proteins, CHEBI identifiers for metabolites etc. |

## 5 Methods

### 5.1 Data model for the Disease Maps Project and the MINERVA Platform

The PD map and COVID-19 Disease Map diagrams are constructed based on literature mining following a set of established community guidelines ^8^. These maps represent key molecular mechanisms of Parkinson’s disease and SARS-CoV-2 infection, respectively, and the following response of the organism ^9, 12, 13, 14^. These diagrams rely on systems biology standards: they are represented using the CellDesigner^19^,^23^ format (https://www.celldesigner.org), which is compliant with the Systems Biology Graphical Notation (SBGN) ^24, 25^ and is built on top of the Systems Biology Markup Language (SBML) ^26^. Annotations of mechanisms and elements are stored following the SBML style for annotations (https://sbml.org/documents/specifications/). As such, they use stable identifiers from the MIRIAM Registry ^27^ and Identifier.org, and descriptors from the COMBINE BioModels Qualifiers (https://github.com/combine-org/combine-specifications/blob/main/specifications/qualifiers-1.1.md#model-qualifiers).

In MINERVA, the maps, including their annotations, can be uploaded and downloaded in different standard or standard-compliant formats, including the CellDesigner format ^19^. These operations can be performed through a dedicated API, which allows retrieval of data under the form of files encoded in the standard formats (XML-based) or JSON elements.

### 5.2 RDA FAIR indicators assessment

We performed the FAIRness assessment of the MINERVA Platform using the FAIRplus-template of the RDA FAIR Data Maturity Model indicators^4^. The RDA FAIR Maturity Model evaluates compliance with each FAIR principle through one or more indicators, focusing on “Data”, “Metadata” and their relationships. Each indicator is associated with an impact level (essential, important, or useful) and indicators target project data, associated metadata, and their relationships. Indicators can be scored in different ways, including binary (true or false, with an option to declare indicators as “not applicable” to a given context) and graded on scales of 1 to 3 or 1 to 5. For our purposes, we opted for the binary approach as it reduces reliance on personal judgement. Evaluation scores were computed for i) all indicators, ii) essential indicators only and iii) non-essential indicators. The results were compared among different groups of indicators, with and without non-applicable indicators in each of these three cases.

### 5.3 Data and associated metadata in the MINERVA Platform

The RDA FAIR Maturity Model indicators target the “Data” and “Metadata” components and their relationships. In MINERVA, the biological content of the maps follows a graph-based representation (with nodes, edges, and their properties). Thus, first we defined the encoding attributes of the data and metadata sets in MINERVA, focusing on their interrelationships to map the meaning of “Data”, “Metadata” and their relationships as used in the FAIR assessment template and the elements (nodes, edges, properties). Table 2 below illustrates this mapping.

**Table 2.** Defining the elements of the disease maps in MINERVA considered as the Data and as the Metadata, respectively, for the FAIR assessment.

| FAIR- targeted components | Elements of the map | Examples |
| --- | --- | --- |
| Data | The model given by the set of individual inter-connected biological maps and describing mechanisms at various biological layers | The Neuroinflammation and the LRRK2 activity maps in the PD map, and the Coagulation and the Apoptosis maps etc. in COVID-19 Disease Map.<br>The complete PD map and COVID-19 Disease Map are openly-available at https://pdmap.uni.lu/minerva/ and at https://covid19map.elixir-luxembourg.org/minerva/ , respectively. |
|  | The biological content, (represented following the COMBINE BioModels Qualifiers: https://github.com/combine- | The mutated LRRK2 protein dephosphorylates PTK2 (in the LRRK2 activity diagram in PD map). Another example |
|  | [org/combine-specifications/blob/main/specifications/qualifiers-1.1.md#model-qualifiers](https://www.ebi.ac.uk/ols/ontologies/combine-specifications/blob/main/specifications/qualifiers-1.1.md#model-qualifiers)). | from Apoptosis diagram in COVID-19 Disease Map includes the protein Orf3a regulates the MAPK14 activation. |
|  | The parameters extracted from the literature for specific mechanisms | Reaction kinetics available in the SBML format <sup>26</sup>. |
|  | The graphical representation of the knowledge itself | The PTK2 dephosphorylation catalysed by the mutated LRRK2 protein (in the LRRK2 activity diagram in PD map) is represented as a set of edges (subgraph) connecting the nodes for the phosphorylated PTK2, PTK2 and mutated LRRK2 proteins.<br>Similarly, the regulation of the Orf3a protein for the MAPK14 activation (in the Apoptosis diagram of the COVID-19 Disease Map) is represented as a set of edges (subgraph) connecting the nodes for the MAPK14, MAPK14 active and Orf3a proteins. |
| Metadata | Annotations such as the publication identifiers associated with the evidence of the integrated knowledge, the UniProt ids for proteins, the species type (e.g. human, virus, animal) etc. In terms of persistence of the compound identifiers (as key feature of the FAIR principles), there are persistent ids for the maps, for the provenience (DOI and PubMed ids for publications), for cell model, species types etc. | The PTK2 dephosphorylation catalysed by the mutated LRRK2 protein (in the LRRK2 activity diagram in PD map) annotations are encoded for several elements: a) the relationship itself (the edge) including the id, the type, the provenience; b) the PTK2 protein (node) including the full name, the posttranslational modifications, the synonyms and other known standard ids such as UniProt, Ensembl, Entrez Gene, HGNC, RefSeq; c) similarly for |
|  |  | the LRRK2 protein (node).<br>Similarly, for the Orf3a regulation of the MAPK14 activation (Apoptosis diagram in the COVID-19 Disease Map. |

In the source files of the map, the SBML format combines data (elements, interactions, and their graphical representation) and metadata (annotations of diagrams and of their components). Specifically, the annotations are included inside the elements they annotate. Thus, from the point of view of the FAIR assessment, metadata points only implicitly to the data they annotate, and metadata is removed when data is removed. This relationship has an impact on several FAIR indicators related to the relationship between the Data and Metadata elements, and to the persistence of the metadata once the data is removed. Consequently, in the current context, indicators such as “RDA-F3-01M: Metadata includes the identifier for the data”, “RDA-A1-01M: Metadata contains information to enable the user to get access to the data”, or the “RDA-A2-01M: Metadata is guaranteed to remain available after data is no longer available” were assessed as non-applicable or not fulfilled (see Results).

### 5.4 Iterative and integrative FAIR assessment using the RDA FAIR Data Maturity Model

Specifically for the assessment process, we created a team including members with expertise on FAIR, the Disease Maps Project (PD map and COVID-19 Disease Map), and the MINERVA platform e.g. ^28, 5, 29, 8, 9, 13, 30^. We performed an iterative series of work meetings, followed by dissemination of the preliminary results to dedicated scientific groups within the FAIRplus and the Disease Maps communities. Their feedback has been integrated iteratively.

## 6 Conclusions and future work

MINERVA is a comprehensive computational environment dedicated to biological diagrams in the systems biomedicine and Disease Maps communities. While exemplified here on the PD map and COVID-19 Disease Map projects, the FAIR assessment is specific to the functionality of MINERVA and to the approaches of the disease map development followed in the Disease Maps Community. Therefore, the results can be applied to all disease maps in MINERVA (https://diseasemaps.org/projects/) and improvement of FAIRness in MINERVA will ultimately have a positive impact on the aforementioned communities as a whole. The current degree of FAIRness of MINERVA is already satisfactory and qualifies it as a valuable knowledge resource for disease maps. However, further work on the FAIRification of MINERVA is already scheduled. The gaps have been identified during the FAIR assessment using the RDA indicators and the FAIRplus tool. We have already started communication with the MINERVA development team regarding the support for the RDA-A2-01M indicator on availability of the metadata in case data is removed or no longer available (as this Essential Indicator is not achieved in the current FAIR assessment). In this direction, the MINERVA-Net ^17^ was developed as a registry to store annotations (metadata) of disease maps available in MINERVA, including the PD map and the COVID-19 Disease Map, and to permit search and exchange of this information across various MINERVA instances. Moreover, due to this evaluation, a functionality was implemented to handle licensing information to address the R1.1 indicator.

Moreover, the FAIR assessment of MINERVA (with example on the PD map and the COVID-19 Disease Map) received good feedback from the communities where we disseminated our results. However, we may need to adapt some of the indicators (e.g. their priorities) to represent more specifically the MINERVA functionality, similarly to some ongoing initiatives, such as the EOSC/RDA-funded project on the adaptation of the FAIR indicators for biosimulations in the “COmputational Modeling in BIology NEtwork” (COMBINE: https://co.mbine.org/) community ^22 31^: https://fair-ca-indicators.github.io/. As the application of FAIR indicators targeting software/tools/structured data is gaining momentum, we aim to collaborate to further scientific efforts on FAIRification in computational Systems Biomedicine. We share the results from this work dedicated to FAIR principles in systems biomedicine as an example on promoting open science in Systems Biomedicine.

## 7 Data Availability

The Parkinson’s Disease Map (PD map) and the COVID-19 Disease Map in MINERVA are openlyavailable at https://pdmap.uni.lu/minerva/ and https://covid19map.elixir-luxembourg.org/minerva, respectively. The FAIR assessment of the MINERVA platform and exemplified on the PD map and COVID-19 Disease Map, is provided as an excel file, openly-available in the zenodo platform at https://doi.org/10.5281/zenodo.10910033.

## 8 Code Availability

no code was developed for this work.

## 10 Acknowledgements

Authors would like to acknowledge all contributors involved in the PD map and COVID-19 Disease Map projects. The MINERVA instances are available online via the ELIXIR-LU project. IB and SG acknowledge FAIRplus Fellowship awards from the IMI FAIRplus project (IMI 802750).

## 11 Author contributions

Conceptualisation: IB, DWe, SG, RS, DWa, MO, VS. Methodology: IB, DWe, DWa, MO, VS. Investigation: IB, DWe, AR, ETI, AM, SG, DWa, MO, VS. Formal analysis: IB, DWe, AR, DWa, MO. Initial draft preparation, IB, DWe, ETI, MO, DWa. Review and editing: all authors. Project coordination: RS, DWa, MO, VS. All authors have read and agreed to the final version of the manuscript.

